# Small RNA-sequencing for Analysis of Circulating miRNAs: Benchmark Study

**DOI:** 10.1101/2021.03.27.437345

**Authors:** Peter Androvic, Sarka Benesova, Eva Rohlova, Mikael Kubista, Lukas Valihrach

**Author notes:** Equal contribution (Peter Androvic, Sarka Benesova).

## Abstract

Small RNA-sequencing (RNA-Seq) is being increasingly used for profiling of circulating microRNAs (miRNAs), a new group of promising biomarkers. Unfortunately, small RNA-Seq protocols are prone to biases limiting quantification accuracy, which motivated development of several novel methods. Here, we present comparison of all small RNA-Seq library preparation approaches that are commercially available for quantification of miRNAs in biofluids. Using synthetic and human plasma samples, we compared performance of traditional two-adaptor ligation protocols (Lexogen, Norgen) as well as methods using randomized adaptors (NEXTflex), polyadenylation (SMARTer), circularization (RealSeq), capture probes (EdgeSeq) or unique molecular identifiers, UMIs (QIAseq). Globally, there was no single protocol outperforming others across all metrics. We documented limited overlap of measured miRNA profiles between methods largely owing to protocol-specific biases. We found that methods designed to minimize bias largely differ in their performance and we identified contributing factors. We found that usage of UMIs has rather negligible effect and if designed incorrectly can even introduce spurious results. Together, these results identify strengths and weaknesses of current methods and provide guidelines for applications of small RNA-Seq in biomarker research.

## Introduction

Circulating microRNAs (miRNAs) found in various body fluids are attractive candidates for clinical biomarkers (1). To identify disease-specific miRNAs, small RNA-sequencing (RNA-Seq) has become a method of choice for its high screening capacity, specificity, sensitivity and ability to quantify isomiRs or detect novel miRNAs (2,3). Despite many advantages, small RNA-Seq protocols suffer from several limitations that obscure quantification. The classical protocol for small RNA library preparation employs two sequential ligations of adaptors to the 3’ and 5’ ends of the miRNAs (in this study represented by Norgen, Lexogen and QIAseq). However, serious quantification bias is introduced in this process due to unequal ligation efficiencies, leading to systematic over- and under-estimation of true miRNA levels (4). The effect is particularly pronounced in biofluids, where miRNA concentration and complexity are rather low (5). Recently, three alternative approaches have been developed to improve quantification accuracy. First approach uses adaptors with randomized nucleotides increasing the chance of effective ligation (NEXTflex) (6); second approach is ligation-free and employs poly-adenylation and template switching during reverse transcription (SMARTer), while third approach relies on ligation of a single 3’ adaptor and subsequent circularization (RealSeq) (7). Additional quantification bias may arise during PCR amplification of libraries. To mitigate PCR bias, unique molecular identifiers (UMIs) have been introduced to identify and remove PCR duplicates (employed in QIAseq protocol), but their effectiveness in small RNA-Seq applications is debated (8,9). In addition, EdgeSeq, a platform using hybridization probes and targeted sequencing readout, specifically designed for ease-of-use in clinical setting, is available as an alternative to small RNA-Seq. Previous comparative studies performed on a subsets of available methods revealed vast differences in their performance (7,9–17). However, how current commercial small RNA-Seq methods perform, particularly in challenging setting such as liquid biopsy samples, is not yet established. Here, we present evaluation of seven commercial small RNA-Seq methods representing all currently available technical approaches for library preparation with focus on their performance for miRNA quantification in human plasma.

## Results

Seven commercially available protocols were used to prepare small RNA-Seq libraries in technical duplicates from: i) human plasma; and ii) equimolar mixture of 962 synthetic miRNAs (henceforth called “miRXplore”) (Fig.1 and Suppl.1). Plasma samples were controlled for isolation artefacts and hemolysis (Methods and Suppl.2), small-RNA library fraction corresponding to miRNAs was gel-purified (Figure S1) and samples were sequenced in a single sequencing run to avoid batch effects (except for EdgeSeq). This design allowed for unbiased comparison of protocol performance with biofluids, as well as detailed evaluation of technical biases. All methods showed high within-protocol reproducibility (Suppl.5, Fig.2), in contrast to low between-protocol reproducibility (Fig.1B); demonstrating that substantial, unique technical bias is introduced by each protocol. To evaluate the extent of this bias, we quantified the log_2_-fold deviation of measured value from expected value for each miRNA in miRXplore sample, where ground truth is known (Fig.1C). EdgeSeq and SMARTer had least bias, while Norgen and Lexogen were most biased, with measured miRNA levels spanning several orders of magnitude (Suppl.4). Surprisingly, single-molecule ligation and circularization approach (RealSeq), recently claiming to significantly reduce bias (7), showed only 21% unbiased miRNAs. In addition, the sequence bias was not reproducible between protocols (Fig.1B, miRXplore), showing that miRNA profiles obtained with different protocols are not comparable.

**Fig. 1.**
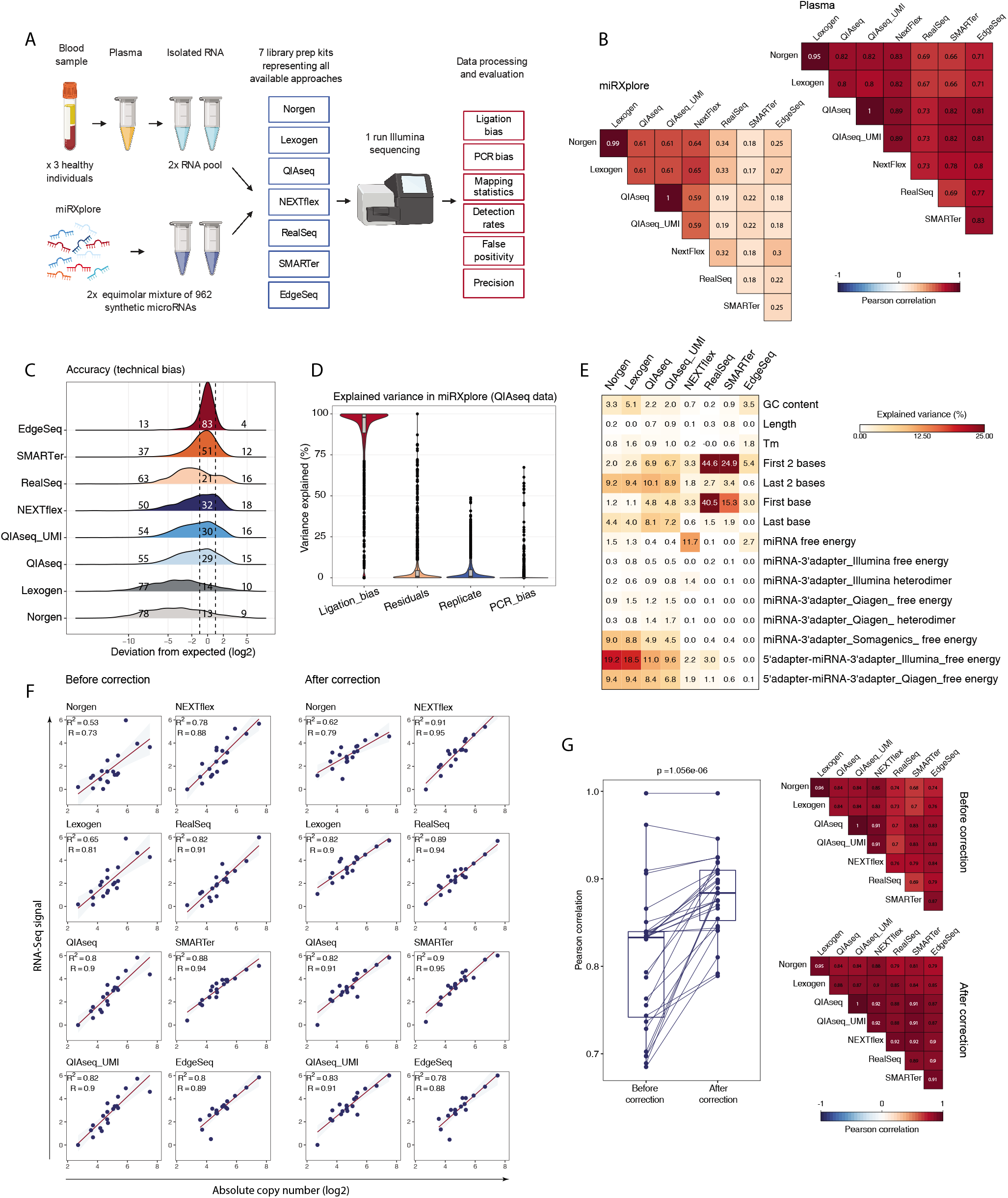
Experimental design, accuracy and technical biases. (A) Schematic representation of study design. (B) Correlation heatmaps showing between-protocol reproducibility for miRXplore and plasma samples. (C) Accuracy determined on miRXplore sample. Density plots show distribution of log_2_-fold change between measured and expected value. Dashed lines show two-fold deviation from expected value; numbers indicate percentage of miRNAs within and outside the two-fold range. (D) Percentage of variance in in QIAseq data (miRXplore sample) explained by ligation bias, PCR bias or replicates. (E) Percentage variance in in QIAseq data (miRXplore sample) explained by miRNA sequence characteristics. (F) Correlation of small RNA-seq with RT-qPCR data measured in plasma before and after data correction with bias ratios learnt from miRXplore samples. (G) Between-protocol reproducibility for plasma samples before and after data correction using bias ratios learnt on miRXplore samples. P-value from two-tailed paired t-test. QIAseq UMI represents data after deduplication, whereas QIAseq means non-deduplicated data.

**Fig. 2.**
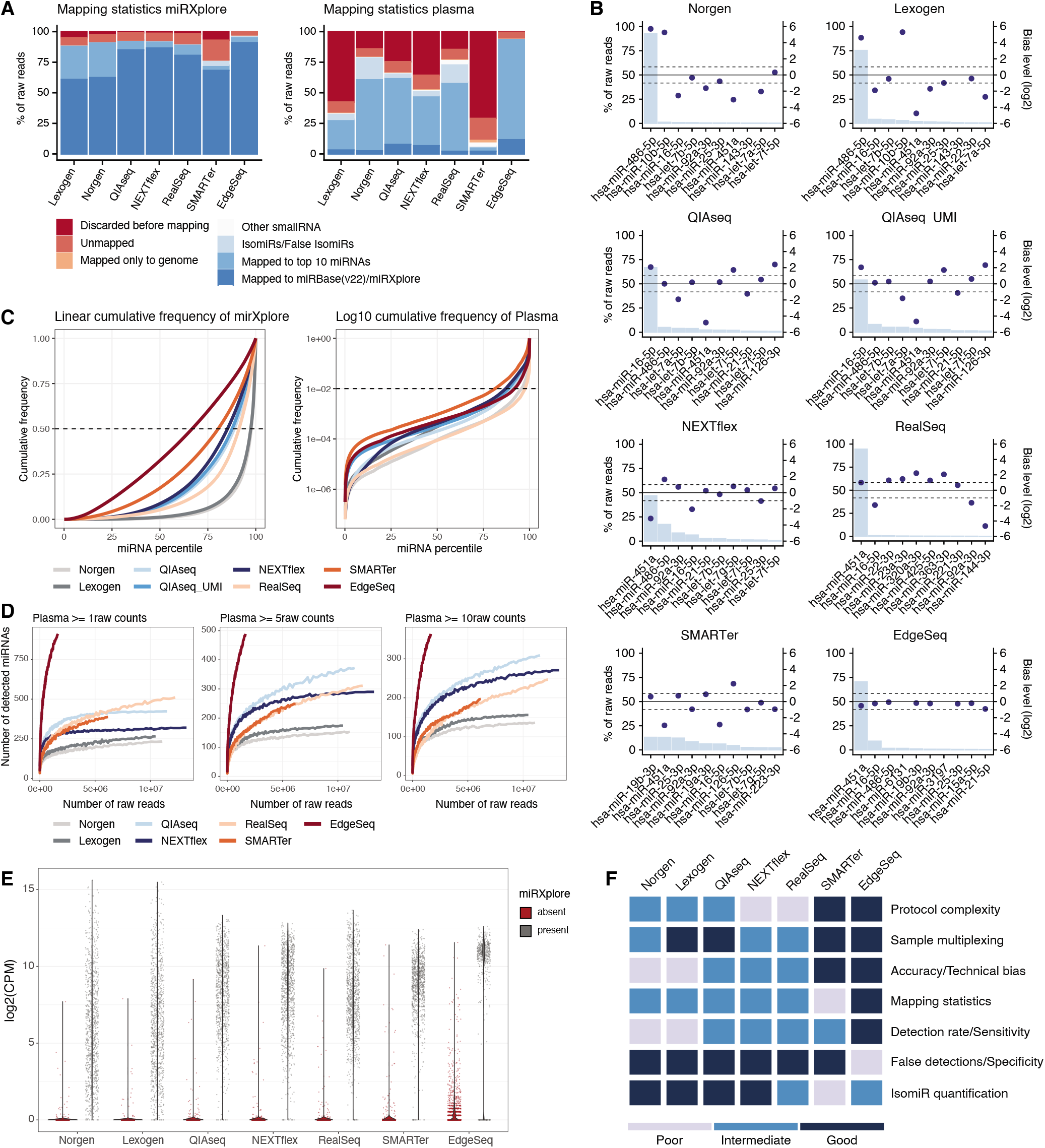
Mapping statistics, sensitivity, false positivity and performance evaluation. (A) Mapping statistics for miRXplore and plasma samples. (B) Top 10 most abundant miRNAs in plasma. Bars (left y-axis) show fraction of raw reads in plasma and dots (right y-axis) show log_2_-fold level of technical bias in miRXplore sample. Dashed lines mark two-fold deviation from expected value. (C) Cumulative frequency of miRXplore and plasma samples in linear and log scale, respectively. Dashed lines indicate cumulative frequency of 50% and 1%. (D) Dependency of number of detected miRNAs in plasma on sequencing depth and various detection thresholds (1, 5, 10 raw reads). (E) Violin plots showing measured level of true and false positive miRNAs measured in miRXplore samples. (F) Final evaluation metrics, for details see Table S4

While previous studies attributed large proportion of the bias to adaptor ligation (4), contribution of PCR to overall bias is often debated, with reports of negligible (4,18) or substantial effect (8,9). In miRXplore sample, we quantitatively evaluated contribution of various factors to overall bias using QIAseq data, which employ UMIs and thus allow separation of PCR contribution from other effects. Ligation bias was highly explanatory for variability in most miRNAs, while PCR bias was overall negligible (Fig.1D). This is in agreement with our previous result showing ligation-free protocols (EdgeSeq and SMARTer) are least biased while ligation-based protocols are most biased overall (Fig.1C). Of note, we identified that short UMI length resulting in insufficient complexity of available UMIs can lead to erroneous overestimation of PCR bias, a likely cause for the misidentification of its contribution in the previous study (Suppl.5, Fig.3). To provide insights into mechanisms leading to biased measurements, we evaluated how miRNA properties explain measured miRXplore values (Fig.1E). First nucleotide in miRNA sequence was highly influential for RealSeq and SMARTer, explaining as much as 44% and 25% of variability. In addition, the identity of the last nucleotides and free energy of adaptor-miRNA construct, but not the miRNA itself, had impact with ligation-based protocols using two defined adaptors including Lexogen, Norgen and QIAseq. Overall, these results demonstrate that ligation, but not PCR is a major source of quantification bias in small RNA-Seq data, and is influenced by complex and technology-specific factors.

Our data revealed that each miRNA is burdened by bias that is specific for each protocol. However, these results were based on balanced mixture of concentrated synthetic miRNAs that may not be fully representative of biological samples such as biofluids, where miRNA concentrations vary broadly, and sequence complexity is lower. To identify how measurements in real biofluid samples are influenced by bias, we quantified absolute abundance of 19 miRNAs in plasma by RT-qPCR (Suppl.5, Fig.4) and correlated it to measured RNA-Seq values (Fig.1F, left). All protocols showed positive correlations with R2 values between 0.53 (Norgen) and 0.88 (SMARTer), although precision for individual miRNAs was often low. In agreement with miRXplore data, Lexogen and Norgen performed worst in this metric. The analysis demonstrates that globally, across-miRNA correlations are relatively preserved in RNA-Seq output from biofluids, i.e., highly abundant miRNAs give high-count values and vice versa. However, values for individual miRNAs are biased and cannot be readily transformed to absolute abundance, making between-miRNA comparisons difficult. We therefore explored if protocol-specific biases learnt from synthetic sample (miRXplore) could be leveraged to correct bias in RNA-Seq data from plasma post-hoc. Indeed, computational correction increased both, correlation of RNA-Seq values with known absolute concentrations (Fig.1F, right) as well as inter-protocol correlation (Fig.1G). These results suggest that protocol-specific biases are preserved (at least to a degree) even between vastly different samples such as plasma and miRXplore. Once learned on the sample with known ground truth, they can be leveraged to both, improve precision of RNA-Seq values and agreement between protocols, potentially facilitating comparisons across studies.

An important decision that researchers face when designing small RNA-Seq experiments is the targeted sequencing depth, which affects the detection rates and cost-efficiency of the experiment. The required sequencing depth is influenced by the ability of protocol to capture molecules of interest and by the proportion of artefact reads. To assess capture efficiency, we evaluated mapping statistics for each protocol (Fig.2A). Note that adaptor-dimers were removed during library preparation in this study and therefore were not mapped (Suppl.5, Fig.1). Whereas the mapping statistics were comparable between protocols with miRXplore, the results revealed substantial differences with plasma samples. The most striking was low mapping rate to miRNAs for SMARTer, which was mostly due to inappropriate read length (Suppl.5, Fig.5). In contrast, targeted approach Edge-Seq showed highest mapping rate of 95%. Importantly, with all protocols the majority of miRNA-mapping reads was consumed by the few highest-ranking miRNAs (Fig.2B,C). As this may reflect true miRNA abundance, but also may be a consequence of bias, we plotted values of the ten most abundant miRNAs in plasma and their corresponding level of bias measured in miRXplore (Fig.2B). In each protocol, except SMARTer and NEXTflex, there was always a single miRNA consuming more than 50% of all mapped reads. Ranks and identity of top 10 miRNAs differed between protocols. Although some miRNAs, such as erythrocyte-specific miR-451 and miR-16 ranked among the highest with all protocols (in agreement with their true abundance, Suppl.5, Fig.4), other miRNAs, such as miR-10b with Norgen and Lexogen appeared to be strongly overestimated (up to 64x) due to bias. To assess allocation of sequencing reads on full miRNA spectrum we further examined curves of cumulative frequencies (Fig.2C). Fast increase in cumulative frequency indicates that even low-ranking miRNAs contribute significantly to the total counts. In miRXplore, number of miRNAs at cumulative frequency of 50% (CF50) would ideally be around 481 (half of 962 miRNAs consumes half of the reads; lower values are better). In agreement with the percentage of unbiased miRNAs, EdgeSeq and SMARTer showed best performance, while Norgen and Lexogen were worst in this metric. In plasma, the shape of ideal curve cannot be known, however it is vastly apparent that the majority of the reads are consumed by few miRNAs. Together, our results show that highly skewed miRNA distribution in plasma is caused by natural miRNA abundance as well as artificial protocol-specific biases and both factors need to be considered to select optimal sequencing strategy.

Considering strong quantification bias of some miRNAs, binary evaluation of miRNA profiles (present/absent) may represent an alternative, more robust approach to identify candidate biomarkers. To characterize variables influencing such analysis, we examined miRNA detection rates at various sequencing depths and count thresholds for each protocol (Fig.2D). While the most of the untargeted protocols approached saturation at 5 million reads, SMARTer and RealSeq further benefited from increased depth. EdgeSeq, QIAseq and NEXTflex detected highest number of miRNAs while Lexogen and Norgen detected fewest. Relative differences between protocols were most pronounced with higher detection thresholds and were retained at various sequencing depths. Interestingly, EdgeSeq detected up to hundreds more miRNAs than any other protocol (Fig.2D). This can be attributed to EdgeSeq high mapping rate (Fig.2A), but it can be also consequence of lower specificity of hybridization probes (19). To investigate this, we plotted measured values for human miRNAs that are present (i.e. true positives) vs human miRNAs that are absent (i.e. false positives) in miRXplore sample (Fig.2E). Indeed, EdgeSeq showed higher false positive rate and higher false signal intensities compared to other protocols, suggesting that its higher detection rate in plasma may be partly due to false positivity. Further, we assessed if miRNAs that were uniquely detected by each protocol in plasma (i.e. miRNAs not detected by any other protocol) are significantly enriched with false-positive miRNAs from miRXplore (Suppl.3). This was indeed the case for EdgeSeq, but not other protocols at all examined detection thresholds. Sequence similarity analysis revealed that false-positive miRNAs detected by EdgeSeq were only modestly similar to true positive miRNAs (Suppl.5, Fig.6), suggesting that false-positive detections may result from incomplete digestion of unbound capture probes, in addition to cross-hybridization. Since miRNA analysis on the level of miRNA variants, isomiRs, is getting more attention in miRNA biomarker studies (20–22), we evaluated the levels of false isomiR detection using miRXplore sample. SMARTer generated most false isomiRs-over 4% of all raw reads, compared to less than 0.4% for other sequencing-based protocols (Fig.2A). Detailed analysis revealed protocol-specific bias between of 3’ and 5’ isomiRs as well as base preferences (Suppl.5,Fig.7,A-B). Whereas some were expected (dominance of 3’ isomiRs with added adenines in SMARTer), prevalence of 3’ isomiRs in EdgeSeq or preference for 3’ thymine addition in RealSeq were unexpected. This raises questions on reliability of isomiRs quantification and warrants careful validation of such data. To sum up, we observed large differences in miRNA detection rate between protocols as well as varying contribution of false positives. Although EdgeSeq captured highest number of miRNAs, it suffered from highest false-positive rate, particularly for miRNAs with low values. Overall, the results suggest caution about spurious detections and highlight the need for data validation by independent technology.

## Discussion

In this study, we compared the performance of all currently available technical approaches for RNA-seq based miRNA analysis in biofluids using a complex set of parameters, including not only data-driven characteristics, but also practical features as protocol complexity or level of multiplexing (Suppl.4). There was no protocol that would stand out as the best across all metrics (Fig.2F). In agreement with other studies (9,10,15), we show that data generated by ligation-free protocols were the least biased, suggesting they may be preferable when quantification of true miRNA abundance is of interest. Particularly, EdgeSeq outperformed others in accuracy, but also in high mapping and detection rate. Other advantages of this platform are automatization minimizing hands-on time and possibility to analyze crude biofluid samples. Although here we analyzed isolated RNA for consistency reasons, Godoy et al.(15) found no major differences between crude and isolated samples. EdgeSeq disadvantages are represented by higher costs of analysis, possibility to quantify only predefined sets of miRNAs and lower specificity, which is in agreement with results of Godoy et al. (15). SMARTer was the second most accurate and the least laborious method from wet-lab perspective. However, its performance was negatively affected by the lowest mapping rate to miRNAs and highest production of artefact reads and false isomiRs, in accordance with previous studies (9,10). On the other hand, SMARTer may be well suited for simultaneous analysis of various classes of small RNAs in a single experiment. Surprisingly, the most recent bias-mitigating approach RealSeq showed accuracy levels similar to NEXTflex and QIAseq, in contrast to results of Barberan-Soler et al.(7) which reported superior accuracy over 70%. Here, we found that circularization approach is not exempt of bias. Considering that RealSeq employs two adaptor ligation steps (one inter- and one intra-molecular) our result seems to be in line with observations that ligation is the most prominent source of bias (5,23). Random adaptors used in NEXTflex represented the third approach in our comparison developed for reduction of ligation bias. In agreement with recent studies (11,12), NEXTflex showed good to average performance in the most of the tested parameters and may be therefore recommended for routine applications in various experimental settings. Lastly, we tested three representatives of traditional ligation-based methods (Lexogen, Norgen, and QIAseq). As expected, Lexogen and Norgen did not perform well in the majority of tested parameters, which is in agreement with the recent literature (11,17). Strong ligation bias leads to misbalanced miRNA profiles, low coverage of the majority of the miRNAs, lower detection rates and therefore to need for higher sequencing depth. Surprisingly, QIAseq that also employs ligation of two defined adaptors, ranked together with NEXTflex among the best in most metrics. We show that this is not due to the usage of UMIs, and since details of the protocol are proprietary, we can only speculate if proper optimization or other bias-mitigating measures are responsible for improved results of QIAseq over Lexogen and Norgen.

Beside the protocol comparison, our data identified several opportunities for improvement of small RNA-Seq analysis in biofluids. Firstly, we documented highly misbalanced miRNA profiles in plasma, where few highly abundant miRNAs consumed most reads (partly due to biological, but also due to technical reasons). New generation of library preparation protocols would therefore benefit from blocking or depleting highly abundant miRNAs such as miR-451 and miR-16. Similar approach was demonstrated on tRNA-halves and improved miRNA detection in serum (24). Secondly, we demonstrated that bias can be learned on synthetic samples with known ground truth and subsequently transferred to improve precision and between-protocol correlation of values in real biofluid sample. Development of advanced computational correction models allowing for complex cross-study comparisons would therefore dramatically increase the utility of publicly available datasets and lead to increase of current knowledge on miRNA profiles in different pathological states. Lastly, contrary to recent reports (8,9), our results suggest that UMIs are superfluous for miRNA quantification and can even lead to serious quantification errors if designed improperly (e.g. with insufficient length). However, our data are based on balanced synthetic template and sample- and protocol-specific factors may pronounce UMIs importance, which needs to be addressed in future studies. For now, we advocate for the developments primarily focused on overcoming ligation bias and improving sensitivity.

Overall, this study can serve as a point of reference for an informed selection of small RNA-Seq method and provide a framework for future development of library preparation protocols and computational methods.

## Methods

### Samples and RNA isolation

Informed consent was obtained from all volunteers participating in the study. All procedures involving the use of human samples were performed in accordance with the ethical standards of Institute of Biotechnology of the Czech Academy of Sciences, and with the Declaration of Helsinki. Blood samples were collected from three healthy volunteers into K_2_ EDTA BD Vacutainer tubes (Beckman Dickinson) and centrifuged within 30 min from collection at 1500 x g for 15 min at room temperature. Plasma fraction was aspirated and transferred into 2 ml tubes (Eppendorf) and centrifuged again for 15 min at 3000 x g. The supernatant was transferred into new 2 ml tubes and stored at −80°C until analysis. Levels of hemolysis were assessed in each sample by measuring absorbance at 414 nm using NanoDrop 2000 (Thermo Fisher) and molecular markers of hemolysis (Suppl.2)(25). Total RNA was isolated starting from plasma aliquots of 250 μl using miRNeasy Serum/Plasma Advanced Kit (Qiagen) according to manufacturers instructions and eluted into 20 μl of nuclease-free water. 1 μl of isolation spike-in mix and 1 μl of GlycoBlue Coprecipitant (Invitrogen) were added at the lysis step as described in (25). Each RNA eluate was assessed for quality of isolation, levels of hemolysis and presence of inhibitors by Two-tailed RT-qPCR panel, as described in (25). RNA eluates were then pooled together to produce standard plasma RNA sample used through the study. An equimolar mixture of 962 synthetic microRNAs (miRXplore Universal Reference) was purchased from Miltenyi Biotec.

### Library Preparation

Libraries were prepared in technical duplicates starting from 5 μl of plasma RNA pool and 5 μl of miRXplore Universal Reference (2×106 copies/μl) according to each manufacturer’s protocol. The version of the protocol, adaptor concentrations and number of PCR cycles for each protocol are listed in Suppl.1. Libraries were quantified on the Qubit 3 fluorometer (ThermoFisher) and Fragment Analyzer (Agilent). Libraries generated by the same protocol were pooled and separated on 5% TBE-PAGE on Mini-PROTEAN tetra cell (BioRad) (Suppl.5, Fig.1). A region representing fragments with RNA inserts of length 22 nt ± 10 nt (i.e. fragments originating from miRNAs) was excised from the gel, DNA was eluted into nuclease-free water and purified with SPRIselect reagent (Beckman Coulter). All libraries were sequenced in one sequencing run on NextSeq 500 high-output (Illumina) with 85 bp single-end reads. 5.8-17.9 million reads per library were obtained with a median of 11 million reads (Suppl.4). EdgeSeq libraries were prepared according to manufacturer’s protocol and sequenced in TATAA Biocenter, Sweden.

### RT-qPCR

Absolute quantification was performed for 35 pre-selected miRNAs using Two-tailed RT-qPCR as described in (26). Briefly, 4 μl of the standard sample (miRXplore) in different concentration (5 to 5xl07 copies/μl) were reverse transcribed using qScript flex cDNA kit (Quantabio) in 20-μl reaction containing pool of miRNA-specific primers. After cDNA synthesis, the total volume was diluted to 200 μl and 2 μl of diluted cDNA were used as a template in 10-μl qPCR reaction containing 1× SYBR Grandmaster Mix (TATAA Biocenter), 0.4 μM forward and reverse primer. The data was processed in Biorad CFX Manager software. Cq values generated by reactions with aberrant melting curves were discarded. For each assay, the standard curves were generated using miRXplore standards and used to calculate absolute miRNA concentration in plasma (Suppl.5, Fig.4). The plasma sample was measured in four technical replicates and two replicates were used for miRXplore standards. After quality control, only 19 miRNAs passing high confidence criteria were used for correlation analysis with RNA-Seq data.

### Data Processing

Raw reads were trimmed with cutadapt tool v1.18 (27) according to the respective library preparation manual. Reads were filtered for length between 15 and 29 bp and subsequently mapped with Bowtie (28) to rRNA and UniVec databases obtained from sortmerna github repository. Reads which did not map to UniVec and rRNA sequences were further mapped to relevant references with STAR (29) using “end-to-end” mode and 5% of sequence was allowed to mismatch. Counting of reads was performed with featureCounts and only uniquely mapping reads were counted. UMI-tools software was used for deduplication before counting of mapped reads in QIAseq samples (30). For comparability with other protocols, non-deduplicated QIAseq data were used for calculation of relevant metrics. Deduplicated QIAseq samples are referred to as “QIAseq_UMI”. Plasma samples were first mapped to human genome (GRCh38.95). Reads mapping to genome were further mapped to mature human miRNA sequences in miRBase v22 (31). Reads which were not mapped to miRBase were further mapped in descended order to isomiRs, tRNA database (435 mature tRNA sequences from gtRNAdb), piRNA database (8 million sequences from piRBase v2) and ncRNA database (36 thousand non-coding sequences from ensemble GRCh38). Mapping to isomiRs and their counting was performed using isomiRROR tool with adjusted settings, when only longer and shorter isomiRs without mismatch in mature sequence were counted. Mapping to other small RNA references was performed with STAR aligner with the same settings as for mapping to miRBase. MiRXplore samples were mapped to miRXplore reference with same settings as plasma samples to miRBase. Raw sequencing data and raw count matrices are available on Gene Expression Omnibus database (https://www.ncbi.nlm.nih.gov/geo/query/acc.cgi?acc=GSE149513) and processed data in Suppl.4. All scripts used for processing data are available on github repository https://github.com/besarka16/Benchmarking-of-small-RNA-seq.

### Evaluation Metrics

If not stated otherwise, all statistics were calculated separately for each technical replicate and their mean values are shown. All samples were normalized by CPM method (divided by total number of reads and multiplied by million). For correlation measures, Pearson coefficients and log_2_-transformed values were used, if not stated otherwise. Technical bias was calculated for each miRNA as a fold change of mean value of two technical replicates from its predicted value. The predicted value was calculated as a number of normalized counts per sample divided by number of miRNAs in miRXplore (962 or 467 for Edge-Seq protocol, respectively). The contribution of PCR bias and ligation bias to overall bias in small RNA-seq was assessed on samples processed by QIAseq protocol with usage of variancePartition R package which employed linear mixed model to separate the variance of multiple variables (PCR bias, ligation bias and technical replicates). Thermodynamic features of miRNAs were calculated by ViennaRNA package 2.0 (32). Contribution of miRNA sequence features to overall bias was assessed using linear model in R with log_2_-fold-deviation as dependent variable. Computational correction of RNA-Seq plasma samples was done using division of normalized counts by ratio of measured and expected expression value in miRXplore sample for corresponding miRNA (33). Dependence of number of detected miRNAs on sequencing depth was assessed by down sampling the raw counts with random generator for binomial distribution in R. The number of miRNAs was used as a number of observations and the number of raw counts belonging to individual miRNAs corresponded to number of trials. The probability of success in each trial corresponded to proportion of raw reads at specific sequencing depth related to the number of raw reads at the original sequencing depth. False positivity was assessed in miRXplore samples, which were re-mapped to human miRNAs (miRBase v22). MiRNAs with ≥ 1 count (in both replicates) and absent from miRXplore reference were considered false hits. Sequence similarity was calculated between all pairs of false hits and miRXplore reference using pairwiseAlignment function from Biostrings R package. Alignment scores were normalized by dividing alignment score by miRNA length and miRNA with maximal score was considered as the best match.

## ACKNOWLEDGEMENTS

This study was supported by Czech Science Foundation P303/18/21942S, Czech Health Research Council NU21-08-00286 and institutional support RVO86652036. We thank vendors and local distributors of small RNA-Seq preparation protocols for discounts or free sample kits that were used in this study.

## AUTHOR CONTRIBUTIONS

P.A. and L.V. designed the study. P.A. prepared standardized material and small RNA libraries. S.B. processed data and performed majority of analyses, with contributions from P.A. E.R. performed RT-qPCR measurements. P.A. and L.V. supervised data analysis. P.A. and S.B. prepared figures and drafted the manuscript. L.V. and M.K. commented on the manuscript and produced the final version. All authors reviewed the manuscript.

---

**Supplementary Note 1: Additional file 1**

Library preparation details, Table S1.xlsx.

**Supplementary Note 2: Additional file 2**

Quality control of standardized plasma samples, Table S2.xlsx.

**Supplementary Note 3: Additional file 3**

Enrichment of protocol-specific miRNAs detected in plasma with false positive miRNAs detected in miRXplore, Table S3.xlsx.

**Supplementary Note 4: Additional file 4**

Complete mapping statistics, raw counts and correlation matrices, ligation bias and metrics table, Table S4.xlsx.

**Supplementary Note 5: Additional file 5**

Supplemental figures, Supplement_figures.pdf.

## Notes

https://github.com/besarka16/Benchmarking-of-small-RNA-seq

https://www.ncbi.nlm.nih.gov/geo/query/acc.cgi?acc=GSE149513

### Competing Interest Statement

The authors have declared no competing interest.

## Bibliography

1. Simone Anfossi, Anna Babayan, Klaus Pantel, and George A Calin. Clinical utility of circulating non-coding RNAs – an update. Nature Reviews Clinical Oncology, 15(9):541–563, sep 2018. ISSN 1759-4774. doi: 10.1038/s41571-018-0035-x.

2. Pauline Chugh and Dirk P. Dittmer. Potential pitfalls in microRNA profiling. Wiley Interdisciplinary Reviews: RNA, 3(5):601–616, 2012. ISSN 17577004. doi: 10.1002/wrna.1120

3. Lukas Valihrach, Peter Androvic, and Mikael Kubista. Circulating miRNA analysis for cancer diagnostics and therapy. Molecular Aspects of Medicine, 72(August): 1–19, 2020. ISSN 18729452. doi: 10.1016/j.mam.2019.10.002.

4. Anitha D. Jayaprakash, Omar Jabado, Brian D. Brown, and Ravi Sachidanandam. Identification and remediation of biases in the activity of RNA ligases in small-RNA deep sequencing. Nucleic Acids Research, 39(21):1–12, 2011. ISSN 03051048. doi: 10.1093/nar/gkr693.

5. Carsten A Raabe, Thean-hock Tang, Juergen Brosius, and Timofey S Rozhdestvensky. Biases in small RNA deep sequencing data. Nucleic Acids Research, 42(3):1414–1426, 2014. doi: 10.1093/nar/gkt1021.

6. Jeanette Baran-Gale, C. Lisa Kurtz, Michael R. Erdos, Christina Sison, Alice Young, Emily E. Fannin, Peter S. Chines, and Praveen Sethupathy. Addressing bias in small RNA library preparation for sequencing: A new protocol recovers microRNAs that evade capture by current methods. Frontiers in Genetics, 6(DEC):1–9, 2015. ISSN 16648021. doi: 10.3389/fgene.2015.00352.

7. Sergio Barberán-Soler, Jenny M Vo, Ryan E Hogans, Anne Dallas, Brian H Johnston, and Sergei A Kazakov. Decreasing miRNA sequencing bias using a single adapter and circularization approach. Genome Biology, 19(1):105, dec 2018. ISSN 1474-760X. doi: 10.1186/s13059-018-1488-z.

8. Yu Fu, Pei Hsuan Wu, Timothy Beane, Phillip D. Zamore, and Zhiping Weng. Elimination of PCR duplicates in RNA-seq and small RNA-seq using unique molecular identifiers. BMC Genomics, (19):531, 2018. doi: 10.1186/s12864-018-4933-1.

9. Carrie Wright, Anandita Rajpurohit, Emily E. Burke, Courtney Williams, Leonardo Collado-Torres, Martha Kimos, Nicholas J. Brandon, Alan J. Cross, Andrew E. Jaffe, Daniel R. Weinberger, and Joo Heon Shin. Comprehensive assessment of multiple biases in small RNA sequencing reveals significant differences in the performance of widely used methods. BMC Genomics, 20(1):513, dec 2019. ISSN 1471-2164. doi: 10.1186/s12864-019-5870-3.

10. Cloelia Dard-Dascot, Delphine Naquin, Yves D’Aubenton-Carafa, Karine Alix, Claude Thermes, and Erwin van Dijk. Systematic comparison of small RNA library preparation protocols for next-generation sequencing. BMC Genomics, 19(1):1–16, 2018. ISSN 14712164. doi: 10.1186/s12864-018-4491-6.

11. Maria D. Giraldez, Ryan M. Spengler, Alton Etheridge, Paula M. Godoy, Andrea J. Barczak, Srimeenakshi Srinivasan, Peter L. De Hoff, Kahraman Tanriverdi, Amanda Courtright, Shulin Lu, Joseph Khoory, Renee Rubio, David Baxter, Tom A.P. Driedonks, Henk P.J. Buermans, Esther N.M. Nolte-’T Hoen, Hui Jiang, Kai Wang, Ionita Ghiran, Yaoyu E. Wang, Kendall Van Keuren-Jensen, Jane E. Freedman, Prescott G. Woodruff, Louise C. Laurent, David J. Erle, David J. Galas, and Muneesh Tewari. Comprehensive multi-center assessment of small RNA-seq methods for quantitative miRNA profiling. Nature Biotechnology, 36(8):746–757, 2018. ISSN 15461696. doi: 10.1038/nbt.4183.

12. Ashish Yeri, Amanda Courtright, Kirsty Danielson, Elizabeth Hutchins, Eric Alsop, Elizabeth Carlson, Michael Hsieh, Olivia Ziegler, Avash Das, Ravi V. Shah, Joel Rozowsky, Saumya Das, and Kendall Van Keuren-Jensen. Evaluation of commercially available small RNASeq library preparation kits using low input RNA. BMC Genomics, 19(1):1—15, 2018. ISSN 14712164. doi: 10.1186/s12864-018-4726-6.

13. Anna M L Coenen-Stass, Iddo Magen, Tony Brooks, Iddo Z Ben-Dov, Linda Greensmith, Eran Hornstein, and Pietro Fratta. Evaluation of methodologies for microRNA biomarker detection by next generation sequencing. RNA biology, 15(8):1133–1145, 2018. ISSN 1555-8584. doi: 10.1080/15476286.2018.1514236.

14. Ryan K.Y. Wong, Meabh MacMahon, Jayne V. Woodside, and David A. Simpson. A comparison of RNA extraction and sequencing protocols for detection of small RNAs in plasma. BMC Genomics, 20(1): 1–12, 2019. ISSN 14712164. doi: 10.1186/s12864-019-5826-7.

15. Paula M. Godoy, Andrea J. Barczak, Peter DeHoff, Srimeenakshi Srinivasan, Alton Etheridge, David Galas, Saumya Das, David J. Erle, and Louise C. Laurent. Comparison of Reproducibility, Accuracy, Sensitivity, and Specificity of miRNA Quantification Platforms. Cell Reports, 29(12): 4212–4222.e5, 2019. ISSN 22111247. doi: 10.1016/j.celrep.2019.11.078.

16. Zachary T. Herbert, Jyothi Thimmapuram, Shaojun Xie, Jamie P. Kershner, Fred W. Kolling, Carol S. Ringelberg, Ashley Leclerc, Yuriy O. Alekseyev, Jun Fan, Jessica W. Podnar, Holly S. Stevenson, Gary Sommerville, Shipra Gupta, Maura Berkeley, Julie Koeman, Anoja Perera, Allison R. Scott, Jennifer K. Grenier, Jeffrey Malik, John M. Ashton, Kara L. Pivarski, Xinkun Wang, Gina Kuffel, Tania E. Mesa, Andrew T. Smith, Jianjun Shen, Yoko Takata, Thomas L. Volkert, Jennifer A. Love, Yanping Zhang, Jun Wang, Xiaoling Xuei, Marie Adams, and Stuart S. Levine. Multisite evaluation of next-generation methods for small RNA quantification. Journal of Biomolecular Techniques, 31(2):47–56, 2020. ISSN 19434731. doi: 10.7171/jbt.20-3102-001.

17. Fatima Heinicke, Xiangfu Zhong, Manuela Zucknick, Johannes Breidenbach, Arvind Y.M. Sundaram, Siri T. Flåm, Magnus Leithaug, Marianne Dalland, Andrew Farmer, Jordana M. Henderson, Melanie A. Hussong, Pamela Moll, Loan Nguyen, Amanda McNulty, Jonathan M. Shaffer, Sabrina Shore, Hoichong Karen Yip, Jana Vitkovska, Simon Rayner, Benedicte A. Lie, and Gregor D. Gilfillan. Systematic assessment of commercially available low-input miRNA library preparation kits. RNA Biology, 17(1):75–86, 2020. ISSN 15558584. doi: 10.1080/15476286.2019.1667741.

18. Ryan T. Fuchs, Zhiyi Sun, Fanglei Zhuang, and G. Brett Robb. Bias in ligation-based small RNA sequencing library construction is determined by adaptor and RNA structure. PLoS ONE, 10(5):1–24, 2015. ISSN 19326203. doi: 10.1371/journal.pone.0126049.

19. Pieter Mestdagh, Nicole Hartmann, Lukas Baeriswyl, Ditte Andreasen, Nathalie Bernard, Caifu Chen, David Cheo, Petula D Andrade, Mike Demayo, Lucas Dennis, Stefaan Derveaux, Yun Feng, Stephanie Fulmer-smentek, Bernhard Gerstmayer, Julia Gouffon, Chris Grimley, Eric Lader, Kathy Y Lee, Shujun Luo, Peter Mouritzen, Aishwarya Narayanan, Sunali Patel, Sabine Peiffer, Silvia Rüberg, Gary Schroth, Dave Schuster, Jonathan M Shaffer, Elliot J Shelton, Scott Silveria, Umberto Ulmanella, Vamsi Veeramachaneni, Frank Staedtler, Thomas Peters, Toumy Guettouche, and Jo Vandesompele. Evaluation of quantitative miRNA expression platforms in the microRNA quality control (miRQC) study. Nature Methods, 11(8):809–815, 2014. doi: 10.1038/nmeth.3014.

20. Danijela Koppers-lalic, Michael Hackenberg, and Renee De Menezes. Non-invasive prostate cancer detection by measuring miRNA variants (isomiRs) in urine extracellular vesicles. Oncotarget, 7(16), 2016. ISSN 1949-2553. doi: 10.18632/oncotarget.8124.

21. Aristeidis G Telonis, Phillipe Loher, Yi Jing, Eric Londin, and Isidore Rigoutsos. Beyond the one-locus-one-miRNA paradigm: microRNA isoforms enable deeper insights into breast cancer heterogeneity. Nucleic Acids Research, 43(19):9158–9175, oct 2015. ISSN 0305-1048. doi: 10.1093/nar/gkv922.

22. Aristeidis G Telonis, Rogan Magee, Phillipe Loher, Inna Chervoneva, Eric Londin, and Isidore Rigoutsos. Knowledge about the presence or absence of miRNA isoforms (isomiRs) can successfully discriminate amongst 32 TCGA cancer types. Nucleic Acids Research, 45(6):2973–2985, apr 2017. ISSN 0305-1048. doi: 10.1093/nar/gkx082.

23. Markus Hafner, Neil Renwick, Miguel Brown, Aleksandra Mihailović, Daniel Holoch, Caroa Lin, John T.G. Pena, Jeffrey D. Nusbaum, Pavel Morozov, Janos Ludwig, Tolulope Ojo, Shujun Luo, Gary Schroth, and Thomas Tuschl. RNA-ligase-dependent biases in miRNA representation in deep-sequenced small RNA cDNA libraries. Rna, 17(9):1697–1712, 2011. ISSN 13558382. doi: 10.1261/rna.2799511.

24. Alan Van Goethem, Nurten Yigit, Celine Everaert, Myrthala Moreno-Smith, Liselot M. Mus, Eveline Barbieri, Frank Speleman, Pieter Mestdagh, Jason Shohet, Tom Van Maerken, and Jo Vandesompele. Depletion of tRNA-halves enables effective small RNA sequencing of low-input murine serum samples. Scientific Reports, 6(October):1–11, 2016. ISSN 20452322. doi: 10.1038/srep37876.

25. Peter Androvic, Nataliya Romanyuk, Lucia Urdzikova-Machova, Eva Rohlova, Mikael Kubista, and Lukas Valihrach. Two-tailed RT-qPCR panel for quality control of circulating microRNA studies. Scientific Reports, 9(1):4255, dec 2019. ISSN 2045-2322. doi: 10.1038/s41598-019-40513-w.

26. Peter Androvic, Lukas Valihrach, Julie Elling, Robert Sjoback, and Mikael Kubista. Two-tailed RT-qPCR: a novel method for highly accurate miRNA quantification. Nucleic Acids Research, 45(15):e144–e144, sep 2017. ISSN 0305-1048. doi: 10.1093/nar/gkx588.

27. Marcel Martin. Cutadapt removes adapter sequences from high-throughput sequencing reads. EMBnet.journal, 17(1):10, may 2011. ISSN 2226-6089. doi: 10.14806/ej.17.1.200.

28. Ben Langmead, Cole Trapnell, Mihai Pop, and Steven L. Salzberg. Ultrafast and memory-efficient alignment of short DNA sequences to the human genome. Genome Biology, 10(3): R25, mar 2009. ISSN 1465-6906. doi: 10.1186/gb-2009-10-3-r25.

29. Alexander Dobin, Carrie A. Davis, Felix Schlesinger, Jorg Drenkow, Chris Zaleski, Sonali Jha, Philippe Batut, Mark Chaisson, and Thomas R. Gingeras. STAR: ultrafast universal RNA-seq aligner. Bioinformatics, 29(1):15–21, jan 2013. ISSN 1460-2059. doi: 10.1093/bioinformatics/bts635.

30. Tom Smith, Andreas Heger, and Ian Sudbery. UMI-tools: Modeling sequencing errors in Unique Molecular Identifiers to improve quantification accuracy. Genome Research, 27(3):491–499, 2017. ISSN 15495469. doi: 10.1101/gr.209601.116.

31. Ana Kozomara, Maria Birgaoanu, and Sam Griffiths-Jones. MiRBase: From microRNA sequences to function. Nucleic Acids Research, 47(D1):D155–D162, 2019. ISSN 13624962. doi: 10.1093/nar/gky1141.

32. Ronny Lorenz, Stephan H. Bernhart, Christian Höner zu Siederdissen, Hakim Tafer, Christoph Flamm, Peter F. Stadler, and Ivo L. Hofacker. ViennaRNA Package 2.0. Algorithms for Molecular Biology, 6(1):26, dec 2011. ISSN 1748-7188. doi: 10.1186/1748-7188-6-26.

33. Anne Baroin-Tourancheau, Yan Jaszczyszyn, Xavier Benigni, and Laurence Amar. Evaluating and Correcting Inherent Bias of microRNA Expression in Illumina Sequencing Analysis. Frontiers in Molecular Biosciences, 6(APR), apr 2019. ISSN 2296-889X. doi: 10.3389/fmolb.2019.00017.

