## Supplemental figures for "Small RNA-sequencing for Analysis of Circulating miRNAs: Benchmark Study"

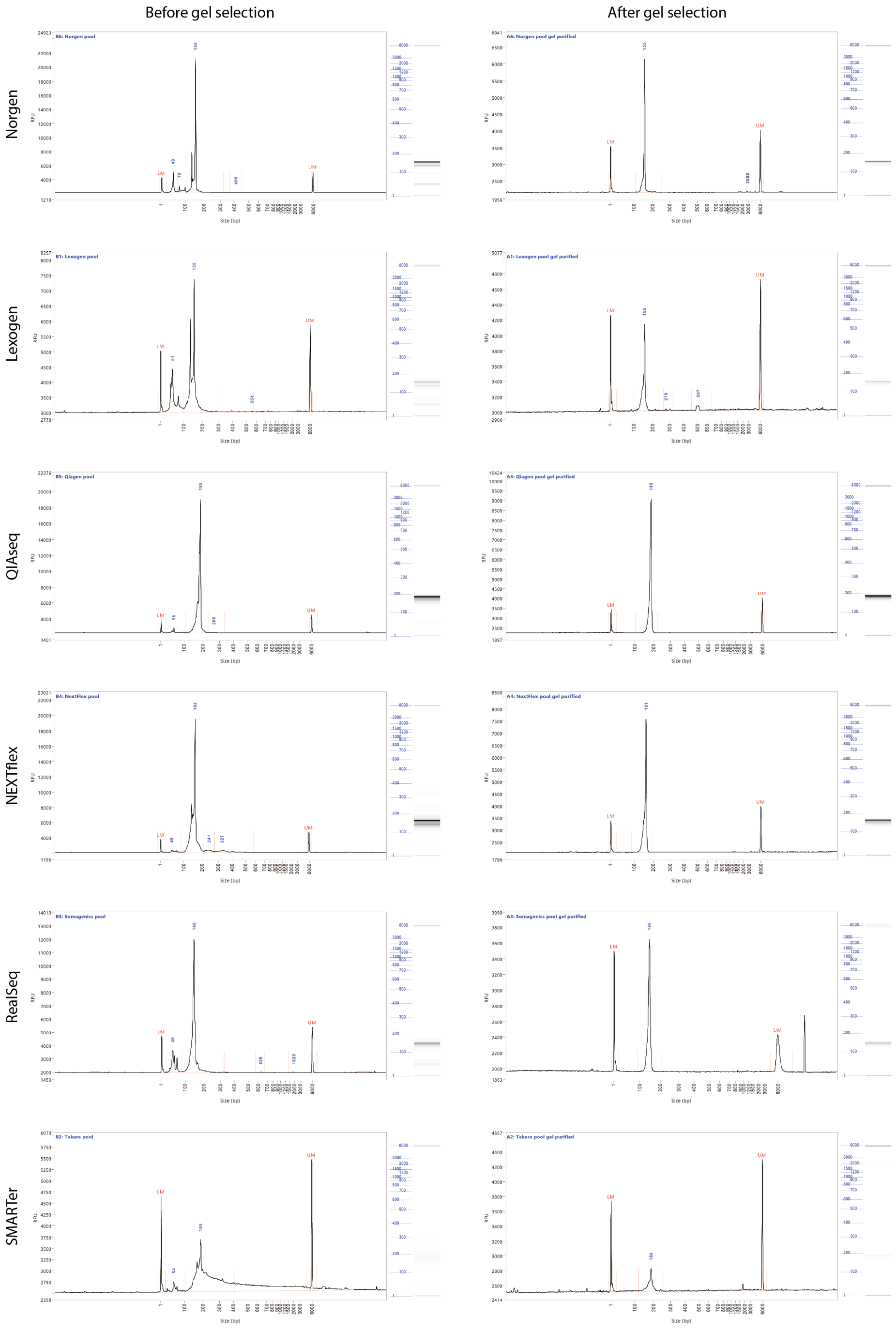


Figure 1: **Electropherograms of libraries measured on Fragment Analyzer.** (A) Pooled libraries before gel selection. (B) Pooled libraries after gel selection.

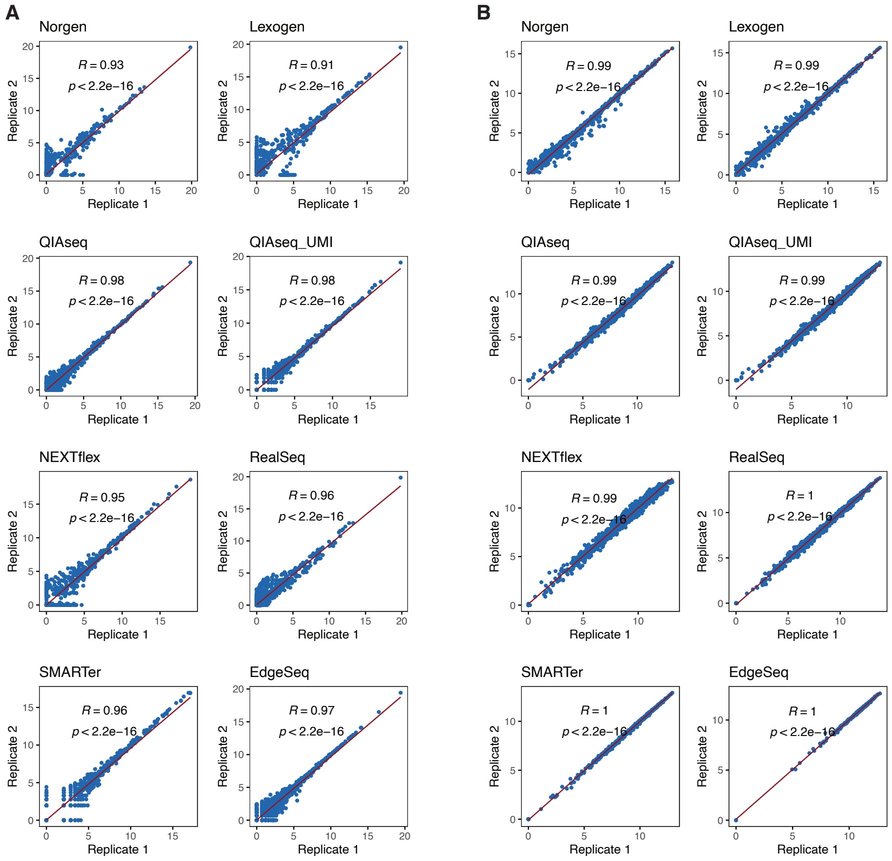


Figure 2: **Within-kit reproducibility.**
(A) Correlation of replicates in plasma samples. (B) Correlation of replicates in miRXplore samples. Values are in log2 scale.

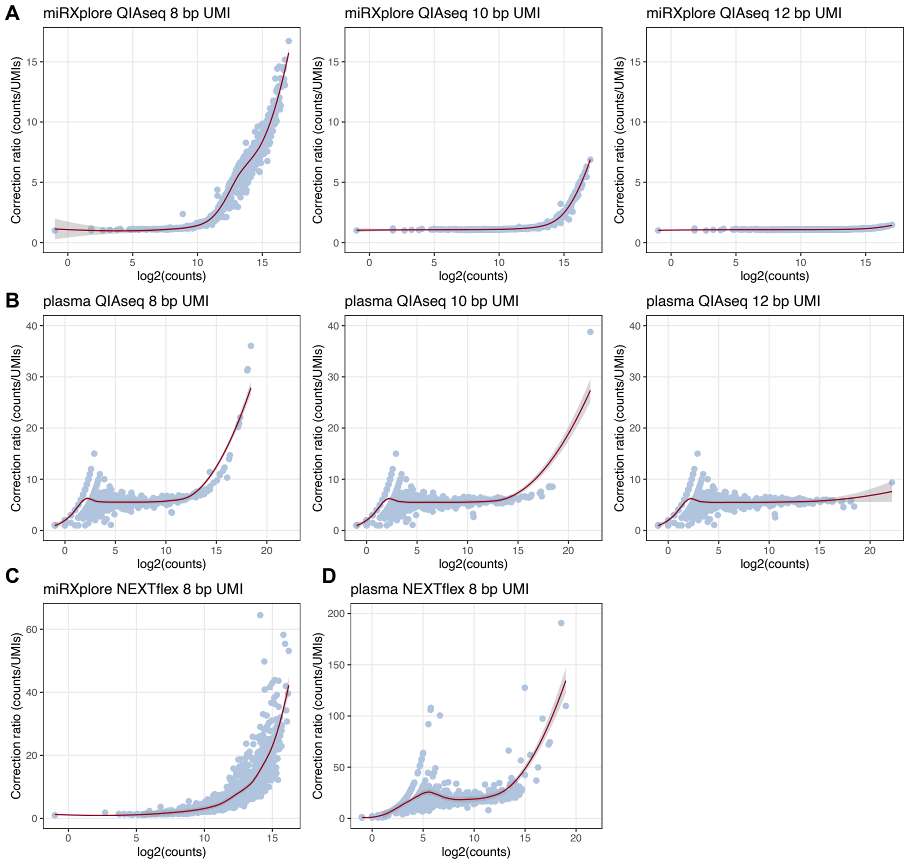


Figure 3: **Insufficient length of UMIs causes erroneous over-correction of miRNA expression values.**

(A) Scatter plots showing uncorrected expression values measured with QIAseq kit in miRXplore sample plotted against UMI correction ratio (raw counts/UMI-corrected counts) when using UMIs with length of 8bp, 10bp and 12bp. Note the sharp increase in correction ratio after miRNA abundance reaches certain threshold. This is caused by insufficient complexity of UMI pool (i.e. two or more miRNA transcripts are bound by the identical UMI, and are erroneously collapsed during computational analysis). This effect is diminished with increasing UMI length. (B) Same as (A), but for plasma sample. Note the similar pattern showing that same scenario applies to real samples. (C-D) Same as (A) and (B), but for NEXTflex kit. Here, we followed the idea of re-purposing random nucleotides adjacent to ligation sites as UMIs, suggested by Wright et al. (2019). We strongly argue against this practice as: i) we show that UMI length of 8 bp is insufficient for standard applications, and ii) placing UMIs at the ligation sites is fundamentally flawed, as it invalidates the assumption of random UMI tagging. Instead, each miRNA species binds the adaptor with the preferred sequence, so in this case more than one miRNA molecule is tagged by the same UMI.

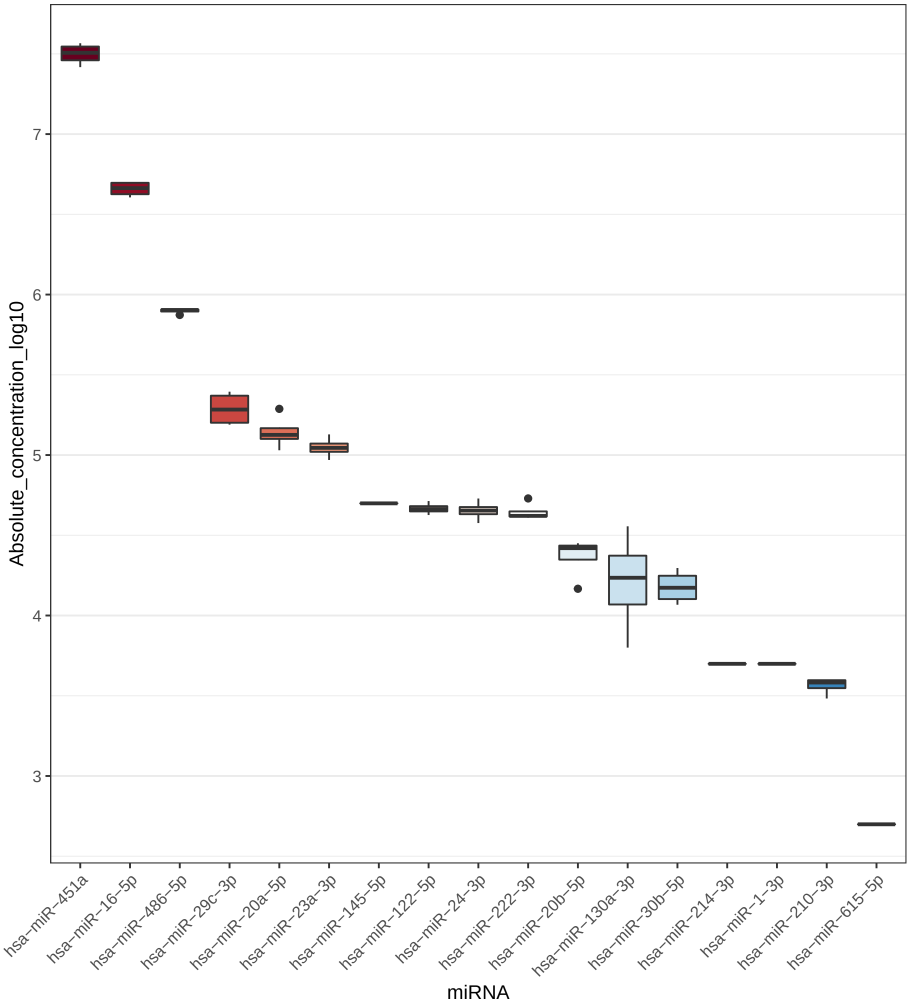


Figure 4: **Absolute concentration of 19 miRNAs measured in plasma samples.** Absolute concentration was determined by RT-qPCR (n=4) using miRXplore as a standard (n=2).

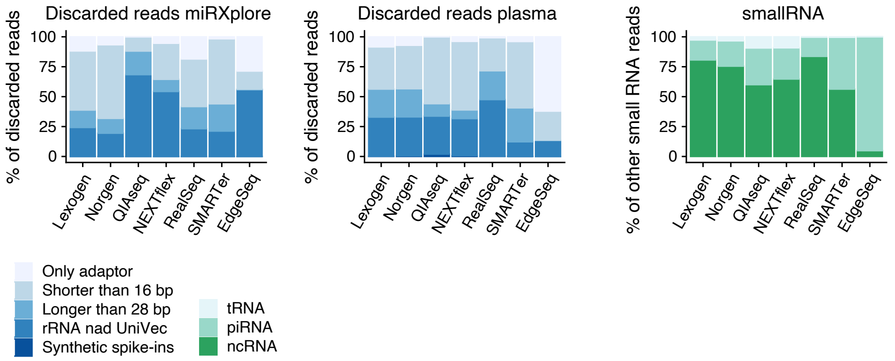


Figure 5: **Mapping statistics of discarded reads and other small RNAs.** (A) Percentage distribution of raw reads discarded before mapping in miRXplore. (B) Percentage distribution of raw reads discarded before mapping in plasma samples. (C) Percentage distribution of raw reads mapped to other small RNA classes than miRNA.

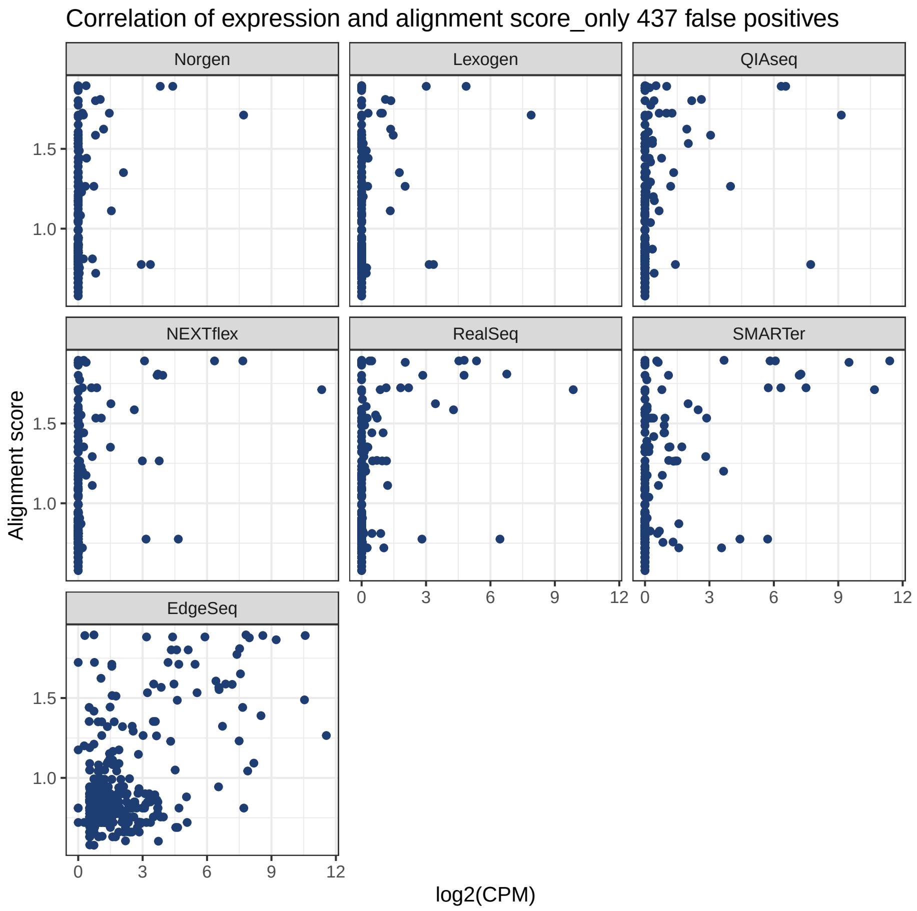


Figure 6: **Sequence similarity analysis.**
Correlation of miRNA measured values in miRXplore samples and alignment score. Scatter plots show 437 miRNAs that were detected as false positive by at least one of the protocols.

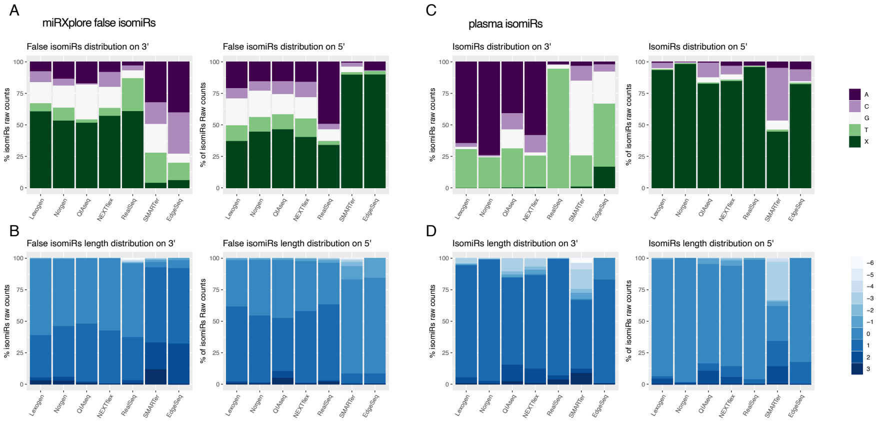


Figure 7: **Mapping statistics of isomiRs.**

(A) Fractions of false isomiR counts in miRXplore samples showing distribution of terminal nucleotides on 3' and 5' end of isomiR. X depicts canonical sequence terminus. (B) Length distribution of false isomiR sequences in miRXplore. Negative values indicate number of missing nucleotides in comparison to canonical sequence, while positive values indicate number of added nucleotides. (C) and (D) are same as (A) and (B), but for all isomiRs in plasma samples.
